# Vagus nerve stimulation using an endovascular electrode array

**DOI:** 10.1101/2023.02.23.529747

**Authors:** Evan N. Nicolai, Jorge Arturo Larco, Sarosh I. Madhani, Samuel J. Asirvatham, Su-youne Chang, Kip A. Ludwig, Luis E. Savastano, Gregory A. Worrell

## Abstract

Vagus nerve stimulation (VNS), which involves a surgical procedure to place electrodes directly on the vagus nerve (VN), is approved clinically for the treatment of epilepsy, depression, and to facilitate rehabilitation in stroke. VNS at surgically implanted electrodes is often limited by activation of motor nerve fibers near and within the VN that cause neck muscle contraction. In this study we investigated endovascular VNS that may allow activation of the VN at locations where the motor nerve fibers are not localized. We used endovascular electrodes within the nearby internal jugular vein (IJV) to electrically stimulate the VN while recording VN compound action potentials and neck muscle motor evoked potentials in an acute intraoperative swine experiment. We show that the stimulation electrode position within the IJV is critical for efficient activation of the VN. We also demonstrate use of fluoroscopy (cone beam CT mode) and ultrasound to determine the position of the endovascular stimulation electrode with respect to the VN and IJV. At the most effective endovascular stimulation locations tested, thresholds for VN activation were several times higher than direct stimulation of the nerve using a cuff electrode; however, this work demonstrates the feasibility of VNS with endovascular electrodes and provides tools to optimize endovascular electrode positions for VNS. The ability to stimulate the VN with endovascular electrodes creates the possibility to stimulate at VN locations that would be otherwise too invasive and at VN locations where structures such as motor nerve fibers do not exist.

## Introduction

Vagus nerve stimulation (VNS) at an electrode surgically placed around the vagus nerve (VN) has been proposed for a variety of conditions including epilepsy, depression, anxiety, obesity, rheumatoid arthritis, heart failure, and more [1-7]. The most surgically accessible location for implanting the VNS electrode cuff is the cervical region as further cranial or caudal would significantly increase invasiveness and risk of complication related to craniotomy or thoracotomy. Despite the relative ease of surgical access and evidence for efficacy, VNS in the cervical region suffers from inconsistent on-target effects, frequent off-target side-effects, and surgical complications.

Inconsistent on-target effects have been observed for several indications of VNS [4,8,9]. One explanation for inconsistent target effects is patient specific anatomical variability. While the gross anatomy of the VN is apparently simple, the internal fascicular organization along the VN is highly variable across both animal models and humans [10,11]. Small stimulation electrodes placed at different locations around the VN in swine and sheep have been shown to elicit different stimulation evoked effects [12,13], suggesting that different groups of fascicles can be selectively activated depending on electrode position. Thus, surgically placed electrodes may be in a sub-optimal location for achieving on-target effect, and cannot be changed without additional surgery. In some applications the ideal stimulation location may be outside the cervical region and cannot be accessed directly without substantially more invasive surgery.

Consistent side effects of VNS in the surgically accessible cervical region are cough, gagging, voice alteration, and other behaviors putatively associated with activation of the laryngeal muscles [4,8,9] caused by electrical stimulation of the motor nerve fibers in the VN [13,14]. These motor nerve fibers have lower stimulation thresholds than the other VN fibers when stimulating with electrodes placed directly on the nerve, and thus can be expected to be recruited by VNS [13-16]. Motor nerve fibers that cause throat muscle contraction are contained within the cervical vagus at every point along the cranial/caudal axis since the motor nerve fibers of the recurrent laryngeal branch traveling from the brain exit the VN within the thorax. Therefore, cervical VNS is limited by activation of motor nerve fibers and subsequent activation of the laryngeal muscles associated with side effects such as cough and gagging. This off-target side effect commonly limits the stimulation current amplitude that can be used [4,8,9].

Surgical complications occur early due to surgery and late due to stimulation and device malfunctions. Early complications include hematoma, infection, and nerve injury that can result in laryngeal dysfunction [17]. Late complications include aforementioned stimulation-evoked activation of the neck muscles, more rarely stimulation-evoked cardiac arrhythmias, and device malfunctions that require additional surgery [4,8,9,17].

Here we investigate a method of VNS from within the adjacent internal jugular vein (IJV). Endovascular VNS has been demonstrated to cause bradycardia in swine, but that prior study did not investigate multiple stimulation configurations on an electrode array, compound nerve action potentials (CAPs), or neck muscle activation as a side effect [18]. Other studies have demonstrated endovascular stimulation of the renal nerve [23], as well as the phrenic nerve [21,22]. Electrical sensing and stimulation of brain tissue using electrodes placed in blood vessels has been demonstrated [19,20]. Our goal was to evaluate endovascular VNS using endovascular electrode arrays and clinically relevant stimulation waveforms and parameters via measurements of evoked CAPs and neck muscle activation [14]. Endovascular VNS has the potential to eliminate all of the aforementioned issues with currently practiced surgical cervical VNS by moving the stimulation electrode in cranial-caudal and axial directions within veins to optimally target multiple areas of the VN without additional dissection, placement of the electrode at any location that the vein travels including regions near or within the cranium and throughout the thorax, and benefiting from the minimally invasive nature of endovascular placement via intravenous access (Figure 1). Additionally, the off-target side effects from activation of cervical region VN motor fibers could be potentially mitigated by expanding the possible VNS implant locations to parts of the nerve that do not contain motor nerve fibers.

**Figure 1:**
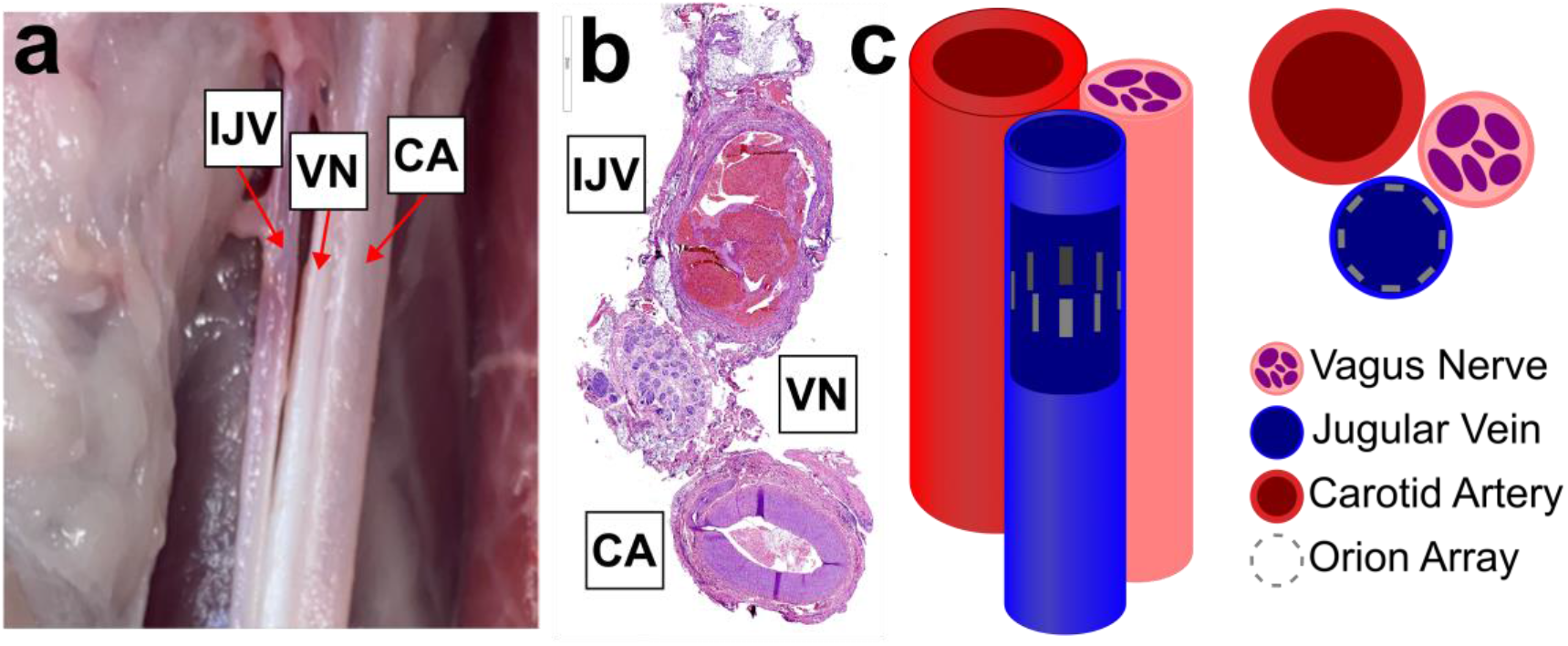
Anatomy and electrode placement. Left, dissection of carotid sheath (sheath removed) showing bundle of internal jugular vein (IJV), vagus nerve (VN), and carotid artery (CA). Middle, histology cross section of swine carotid sheath showing one possible orientation of IJV, VN, and CA. Right, cartoon of the carotid sheath components with a cutout of the IJV showing the Orion electrode array as eight grey rectangles.

Although our vision for endovascular VNS is to stimulate regions of the VN without motor nerve fibers, we performed these initial experiments in the cervical region where motor nerve fibers are within the VN trunk in order to use the evoked muscle responses as another output of VNS to understand how electrode position in the vein affects VNS evoked responses. Here we present stimulation-evoked changes in VN CAPs and neck muscle contraction in response to multiple endovascular electrode configurations. We also demonstrate non-invasive methodologies for targeting the VN location with respect to IJV.

## Methods

### Animals

All animal care, use, and procedures were approved by the Institutional Animal Care and Use Committee (IACUC) of Mayo Clinic. Six male domestic pigs were included in the study weighing 30 ± 1 kg (mean ± standard deviation, SD). All procedures were performed in an acute terminal study lasting between 8 and 10 hours.

### Anesthesia

Pigs were sedated using an intramuscular injection of telazol (6 mg kg^−1^), xylazine (2 mg kg^−1^), and glycopyrrolate (0.006 mg kg^−1^), then intubated and anesthetized using isoflurane (1.5%–3% isoflurane in room air). Analgesia (fentanyl, 5 μg kg^−1^ bolus i.v., followed by 5 μg kg^−1^ h^−1^ per hour) was provided via continuously administered saline (0.9%). Depth of anesthesia was monitored – and isoflurane concentration was adjusted – via changes in heart rate, respiration, and end-tidal CO_2_. Animals were euthanized using pentobarbital (100 mg kg^−1^ i.v.).

### Surgical Procedure

Pigs were positioned supine and femoral vein access was gained under ultrasound guidance, using the Seldinger technique. A control angiogram was performed to confirm vessel patency and then heparin was administered through a cannula in the animal’s ear. Then, under roadmap visualization (Artis Z system Siemens Healthcare, Forchheim, Germany) a soft tip guide wire was introduced into the femoral vein and advanced through the inferior vena cava (IVC), right atrium and superior vena cava (SVC) into the IJV. Angiograms were performed to differentiate the external jugular vein from the IJV and ensure the guide wire was in the IJV (Supplementary Figure 1). An 18Fr sheath (Sentrant, Medtronic, Minneapolis, MN, USA) was then advanced over the guide wire and positioned within the (IJV). Using the sheath as a guide, a cardiac mapping catheter (Orion, Boston Scientific, Marlborough, MA) was then advanced within it and positioned within the IJV in parallel to the VN.

A 2-3 cm incision was made above the thyroid cartilage and custom built stainless steel bipolar needle electrodes were placed into the cricothyroid and cricoarytenoid muscles for measurement of electromyography (EMG).

An incision was made through the skin and superficial fat layers between the mandible and sternal notch using a cautery. The carotid sheath was exposed at a window approximately 1 cm long where custom built longitudinal intrafascicular electrodes (LIFE) could be placed into the VN. This surgical cutdown was approximately 6-8 cm cranial to the location of the Orion in the IJV to avoid disruption of the natural conformance between the IJV and VN.

### Endovascular Stimulation Device

The Orion mapping catheters (Orion RC64S, Boston Scientific, Marlborough, MA) used in this study were purchased from eSutures.com. The impedance of every electrode contact was measured before use in an animal experiment to ensure functionality. An extension wire for the Orion (Umbilical Cable RAUMBILICAL2, Boston Scientific, Marlborough, MA) was modified to connect multiple electrode contacts together to increase electrode surface area for application of higher stimulation amplitudes (See Supplementary Figure 2). This modification resulted in a ring of eight stimulation contacts that lie equally spaced around the circumference of the IJV (Figure 1c).

### Stimulation and Recording Equipment

Electrophysiology recordings were performed on a Neuralynx acquisition system (“Freelynx”, Neuralynx, Bozeman, MT) and sampled continuously at 30kHz (anti-aliasing filter = 8.5 kHz, gain = 192). Stimulation was applied using an A-M Systems Isolated Pulse Stimulator (Model 2100, Sequim, WA).

### Experimental Protocol

The electrical stimulation parameters used in this study were monopolar configuration, symmetric biphasic pulses, cathodic leading, 200 microsecond per phase, variable amplitude ranging from 0.1 mA to 5 mA, 25 Hz repetition rate, pulse trains lasting between 3 and 10 seconds. Test stimulations ranging from 1 to 3 mA were performed to determine the starting position of the Orion stimulation array for the experiment. Briefly, stimulation was applied at each electrode around the circumference of the vein while measuring EMG of the CA and CT muscles. The electrode that produced the lowest EMG threshold was then stimulated at several locations along the length of the vein to determine the final position along the length of the vein and nerve.

Pulse trains of randomized stimulation amplitude – between 0.1 to 5 mA – were performed at each electrode contact while measuring ENG and EMG. After all stimulation amplitudes were performed for every electrode contact in monopolar configuration, stimulation was applied in several bipolar configurations while recording ENG and EMG. A neuromuscular junction blocking agent, vecuronium, was injected to verify identity of EMG motor evoked potentials. Transection of the recurrent laryngeal branch and vagus trunk were performed to verify the identity of motor evoked potentials and compound action potentials, respectively. Signals that were removed by transection of the recurrent laryngeal branch were identified as “RL-mediated” motor evoked potentials (MEPs). Signals that were removed by transection of the vagus trunk were identified as compound action potentials (CAPs). After euthanasia, post-mortem dissection was used to confirm the relative position of the Orion electrode contacts with respect to the VN (Supplementary Figure 3). Whole carotid sheath – VN, IJV, and carotid artery – were collected in some animals for histological analysis.

### Data Analysis

All data analysis was performed in Matlab R2018B (Mathworks, Natick, MA). ENG and EMG signals were filtered and pre-processed as described in our previous report [14]. Briefly, a 400 sample median high-pass filter was applied to the raw 30kHz signal to produce an approximation of the baseline drift, which was then subtracted from the raw signal to remove baseline drift. Stimulation triggered average traces were calculated for each stimulation pulse train.

### Histological Analysis

We collected whole carotid sheath – VN, IJV, and carotid artery – from locations near the placement of the endovascular stimulation electrode and near the surgical cutdown for placement of the recording longitudinal intrafascicular electrodes. Samples were placed in 10% neutral buffered formalin for at least 24 hours, processed according to a standardized protocol, and embedded in paraffin. Each block was sectioned into 5 µm thick slices, which were stained using hematoxylin and eosin (H&E). Slides were imaged using a Motic EasyScan (Emeryville, CA) at 20x magnification.

## Results

### Verification of Electrophysiology and Electromyography Signals

In another report, members of our team demonstrated the use of a neuromuscular junction blocker and nerve transection to verify the sources of signals in electrophysiology (ENG) and electromyography (EMG) recordings [13-14]. Given the indirect nature of endovascular stimulation and that endovascular electrodes are placed “blind” without surgical cutdown, our first objective was to verify the sources of ENG signals evoked by endovascular VNS. Figure 2a shows stimulation triggered average signals recorded at longitudinal intrafascicular electrodes (LIFEs) in the vagus trunk in response to a range of stimulation amplitudes (y-axis). Figure 2b shows that transection of the RLN branch abolishes the ∼6 ms latency signal observed in ENG recordings (as well as EMG recordings, not shown), demonstrating that that signal is caused by neurotransmission along the RLN branch resulting in contraction of the cricoarytenoid neck muscle. The latency of this signal is consistent with previous reports of neck muscle contraction in response to activation of the motor nerve fibers within the VN trunk [13,14], as opposed to activation of those same motor nerve fibers in the RLN branch which would have a shorter latency. Figure 2c shows that transection of the VN trunk between the location of the endovascular stimulation electrode and the LIFE electrodes in the nerve abolishes the ∼1-2 ms latency signals observed in the ENG recordings, demonstrating these signals are caused by neurotransmission along the VN trunk.

**Figure 2:**
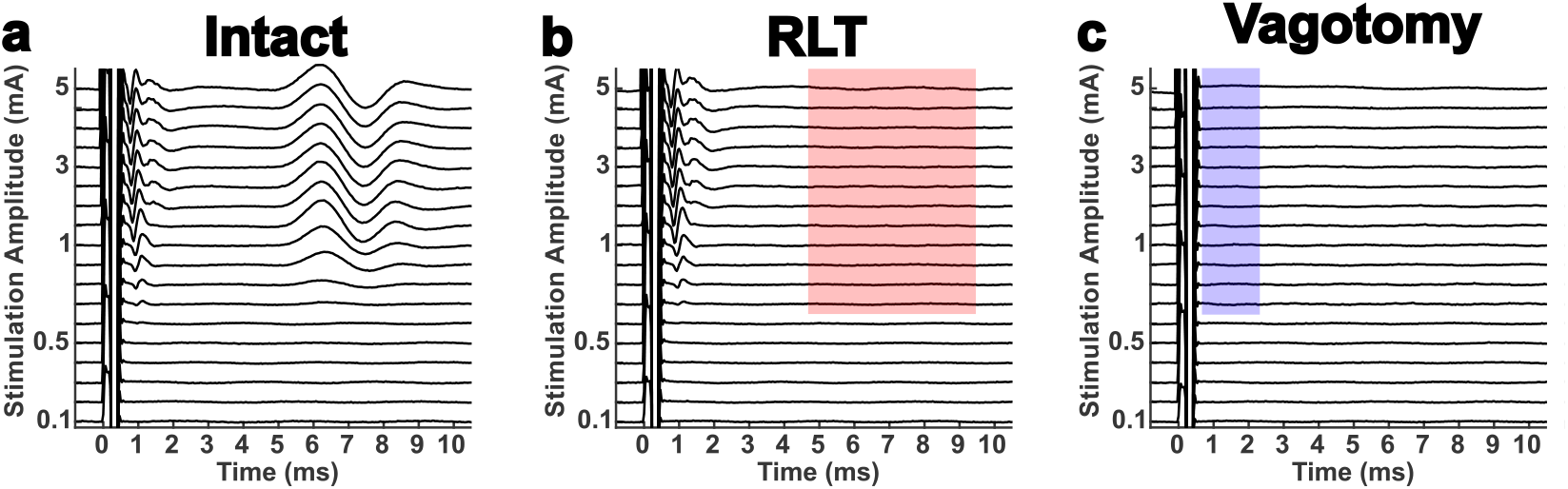
Verification of signal identities via transection of recurrent laryngeal branch (RLT) for motor evoked potentials (MEPs) and transection of the vagus nerve trunk (Vagotomy) for compound action potentials (CAPs). All traces shown are computed from electroneurogram (ENG) recordings made at longitudinal intrafascicular electrodes (LIFEs) sewn into the vagus trunk in response to stimulation at the Orion electrode array within the internal jugular vein. **a)** Baseline, or while the nerve trunk and recurrent laryngeal branch are intact, stimulation triggered median traces in response to randomized presentation of various stimulation amplitudes (y-axis). **b)** Stimulation triggered median traces in response to the same set of stimulation amplitudes after transection of the recurrent laryngeal branch. Red shaded box highlights absence of the signal that was recorded during the intact state, indicating this signal was caused by neurotransmission along the recurrent laryngeal branch and are thus identified as RL-mediated EMG artifacts contaminating the ENG recordings. **c)** Stimulation triggered median traces in response to the same set of stimulation amplitudes after vagotomy. Blue shaded box highlights absence of the signals that were recorded during the intact and RLT states, indicating that these signals were caused by neurotransmission along the vagus trunk and are thus identified as CAPs.

### Activation via different endovascular electrode contacts demonstrates importance of proximity to nerve

Given that stimulation electrical potential is known to decay over space [24], we hypothesized that stimulation contacts on the Orion array that were closer to the VN would have lower activation thresholds for stimulation-evoked effects than the other more distant electrodes (See Figure 1 for Orion array orientation within the jugular vein). There are 8 evenly spaced identical electrodes around the circumference of the modified Orion array, and we first evaluated ENG and EMG responses to monopolar configuration stimulation at each individual electrode. In animals where we observed stimulation evoked responses at most or all of the electrodes (ENG n=3, EMG n=4), we observed an inverted U-curve response profile for the threshold of ENG and EMG signals with respect to electrode position around the vein where one of the electrodes (labeled as 0 degrees) had the lowest threshold (See Figure 3). See Supplementary Figure 4 for a summary of responses across animals.

**Figure 3:**
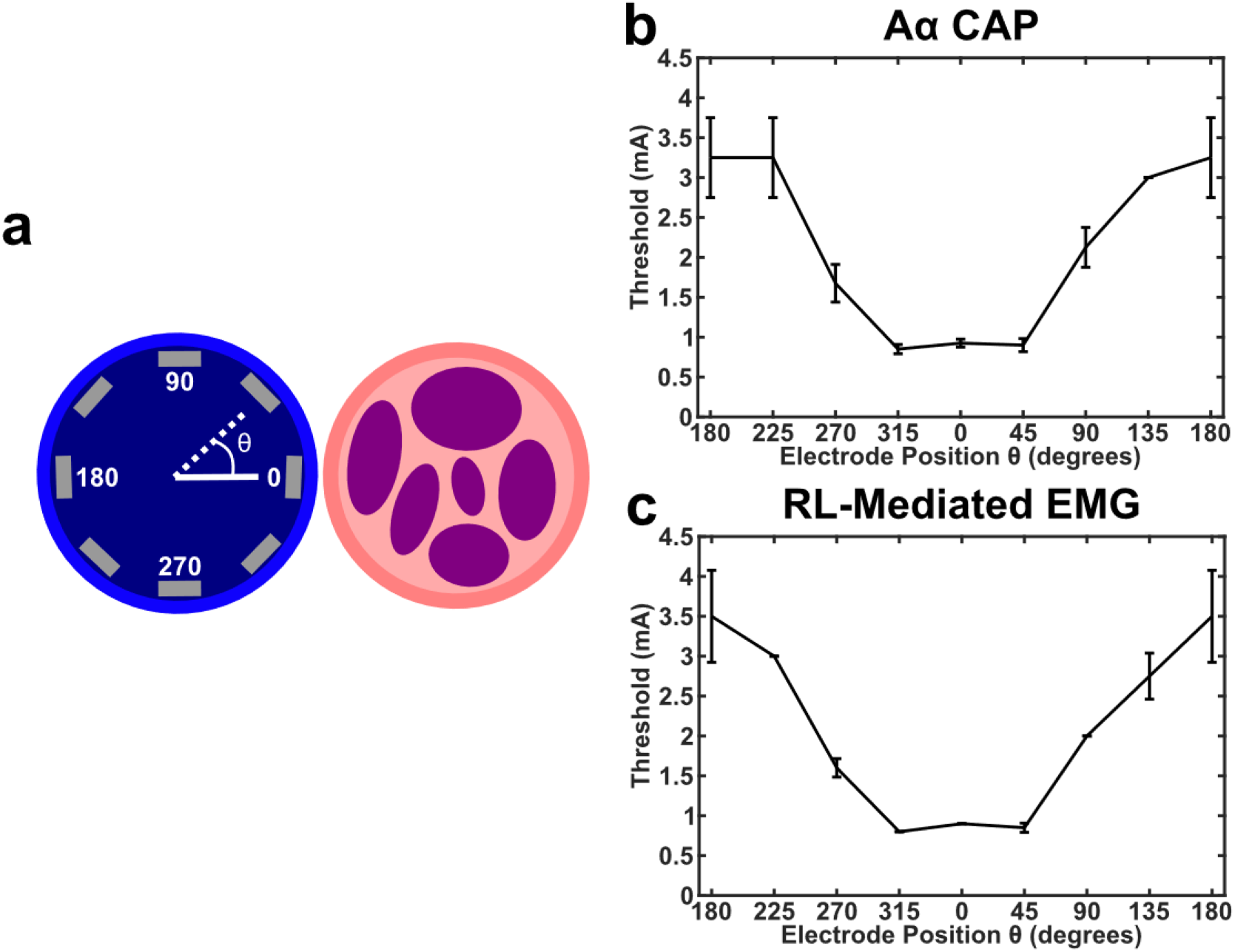
Endovascular circumferential electrode array and VNS activation demonstrating that electrode position within vein is critical for optimal activation of the nerve. **a)** Cartoon shows the internal jugular vein as the blue circle (left), vagus nerve trunk as the pink circle (right), and the 8 electrode contacts as the gray rectangles. Position of stimulation electrode contacts were verified with postmortem dissection, and data from each animal was binned such that the electrode closest to the VN were labeled as θ = 0. **b)** Thresholds for activation of the Aα CAP in response to monopolar stimulation at each of the 8 contacts in one animal (n = 4 LIFE electrodes, black line is mean, error bars are standard deviation). **c)** Same as **b**, but for RL-mediated EMG (n = 2 needle electrodes, black line is mean, error bars are standard deviation).

### Bipolar stimulation configurations

After determining that the position of a monopolar electrode around the circumference of the vein has a clear effect on the threshold of stimulation evoked responses, we investigated the effect of bipolar stimulation configurations. Figure 4a shows the dose response curve for monopolar stimulation at the stimulation electrode contact found to have the lowest evoked CAP threshold, where threshold is approximately 700 µA. Figure 4b shows the dose response curve for bipolar stimulation when the cathode and anode are near each other (See Figure 4, Left Diagrams), where the CAP threshold is approximately 2000 µA and the difference between threshold to saturation is wider than the monopolar configuration. Figure 4c shows the dose response curve for bipolar stimulation when the cathode and anode are far apart, where the CAP threshold is approximately 900µA and difference between threshold to saturation is more similar to monopolar configuration. These differences in threshold for electrode configuration were consistent across animals where monopolar configuration (Figure 4a) showed the lowest evoked thresholds (RL-mediated EMG 960 ± 230 µA mean ± std n = 4 pigs; Aα CAP 1080 ± 550 µA mean ± std n = 3 pigs), bipolar far (Figure 4c) showed the highest evoked thresholds (RL-mediated EMG 2150 ± 470 µA mean ± std n = 4 pigs; Aα CAP 2500 ± 1000 µA mean ± std n = 3 pigs), and bipolar close (Figure 4b) showed something in between (RL-mediated EMG 1080 ± 260 µA mean ± std n = 4 pigs; Aα CAP 1300 ± 430 µA mean ± std n = 3 pigs). A summary of these responses can be found in Supplementary Figure 5.

**Figure 4:**
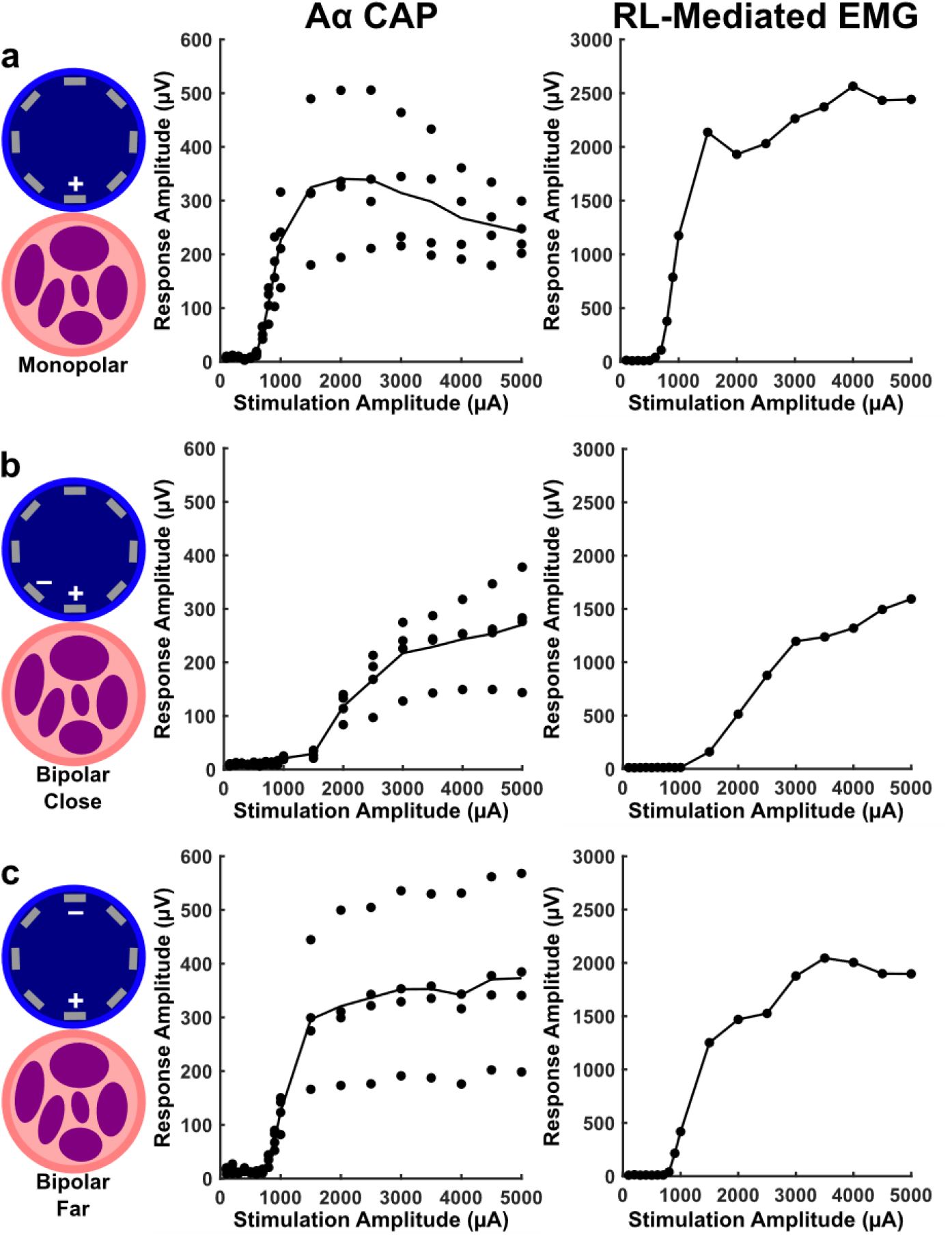
Monopolar vs bipolar stimulation configurations. **a)** Responses to monopolar stimulation where the electrode closest to the nerve is being stimulated (+) and the return electrode is a needle in the left forelimb of the pig. **Left** column is a cartoon illustrating position of the electrodes used for stimulation with respect to the vein and nerve. **Middle** column shows amplitude of Aα CAPs in response to monopolar stimulation in one animal (n = 4 LIFE electrodes, black line is mean). **Right** column shows amplitude of RL-mediated EMG in response to monopolar stimulation in one animal (difference in response to 2 needle electrodes shown). **b)** Same as **a**, but in response to bipolar “far” stimulation where the stimulated electrode (+) has a return at the electrode furthest away marked (-). **c)** Same as **b**, but in response to bipolar “close” where the stimulated electrode (+) has a return at the nearest electrode marked (-). The electrode marked with a (+) was stimulated using a cathodic leading pulse in all configurations.

### Methodology to determine nerve and vein orientation non-invasively during surgery

Given that these studies were performed using ultrasound imaging to obtain vascular access and fluroscopic imaging to guide the Orion catheter to the IJV, we also investigated the use of these imaging technologies to help determine the orientation of the nerve with respect to the vein. Figure 5a shows a B-mode ultrasound (9 MHz center frequency) image through the skin of a pig neck above the carotid sheath with arrows identifying the VN, carotid sheath, and IJV. The vein and artery can be easily identified using doppler imaging which highlights blood flow. The vein and artery can be identified from one another by compressibility, the vein collapses with light pressure from the ultrasound probe. The nerve can be identified by a characteristic “honey-comb” structure, with a white outline and black holes [27]. A 19 gauge needle was placed via ultrasound guidance such that the tip was near the edge of the VN. Immediately after, fluoroscopy was used to guide the Orion electrode to the IJV (see Methods *Surgical Procedure*). Biplane fluoroscopy was used to move the Orion electrode as close to the needle as possible. The fluoroscopy machine was set to cone beam mode, images were collected for one rotation of the beam, and the image in Figure 5b was reconstructed. This demonstrates how existing tools can be used to guide an endovascular electrode array to a location where the VN and IJV are closely opposed.

**Figure 5:**
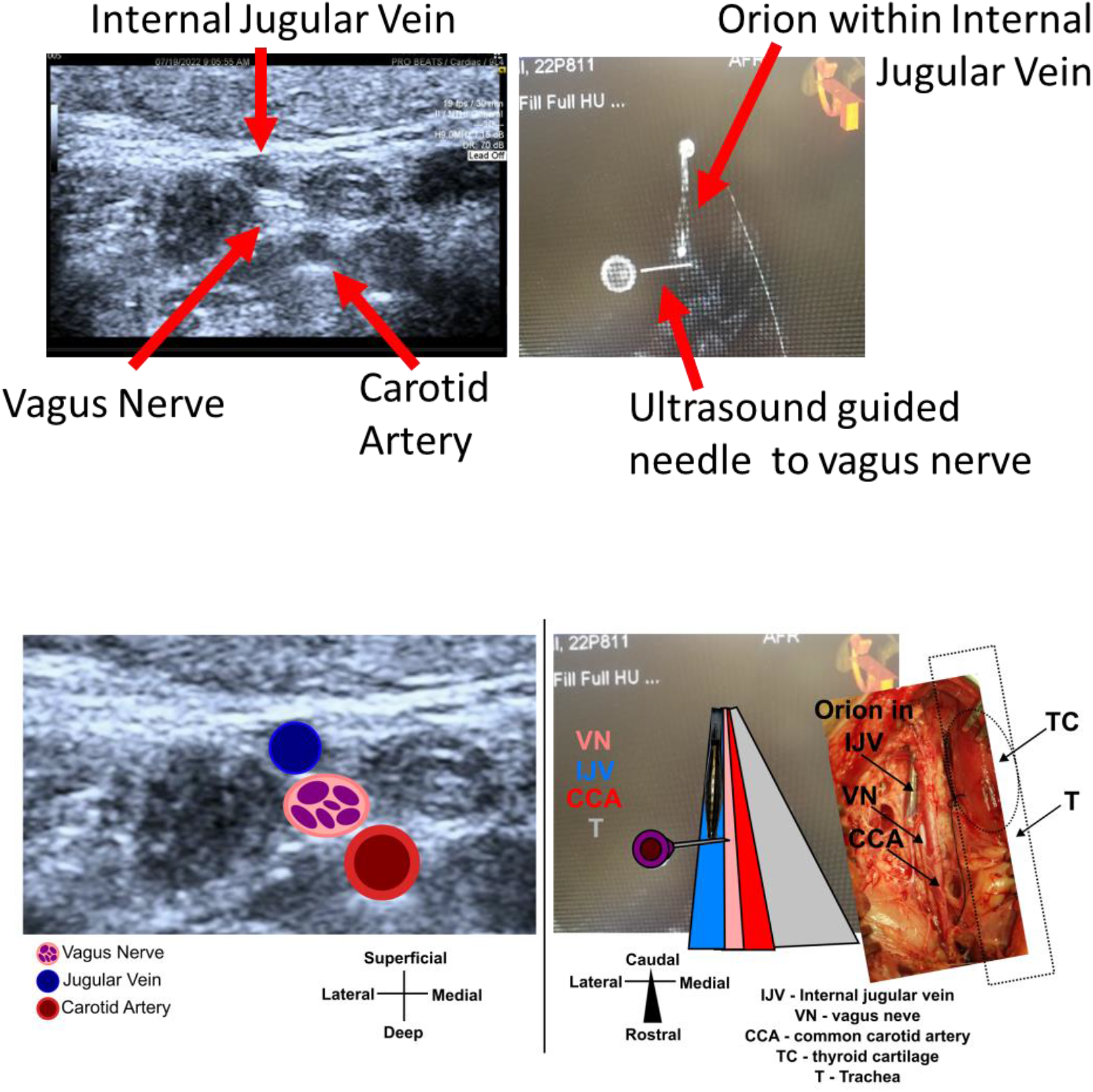
Ultrasound imaging to determine the position of the vagus nerve with respect to the internal jugular vein and cone beam DynaCT to navigate the intravascular electrode to that position. **Top)** Left shows 9 MHz center frequency ultrasound imaging of the cervical region, arrows highlight and label each structure. The internal jugular vein and carotid artery were confirmed via color doppler, and the internal jugular vein was further confirmed via compressibility. The vagus nerve was determined via characteristic “honeycomb” structure. A needle was placed near the vagus nerve via ultrasound guidance. Right shows cone beam DynaCT mode for fluroscopy imaging of the cervical region, arrows highlight the intravascular Orion electrode and the needle placed near the vagus nerve via ultrasound. **Bottom)** Same ultrasound and fluoroscopy images as **Top**, but with cartoons overlaid to help guide readers not familiar with interpreting ultrasound and CT images. Right overlaid photograph (tilted to match orientation of CT image) shows surgical cutdown from this animal which verifies the position of the Orion with respect the vagus nerve.

## Discussion

Electrical stimulation applied within blood vessels is known to affect nearby nervous structures, including the VN [18-23]. The endovascular VNS studies that exist – to our knowledge – have not demonstrated methodology to selectivity target the VN near the vessel, and have not demonstrated direct measures of vagus activation such as evoked nerve compound action potentials. Our goal was to employ an endovascular stimulation electrode array that allows for multiple stimulation configurations and use common clinical parameters of VNS [14,8,9,4] while measuring evoked compound action potentials within the vagus trunk. Additionally, we investigated design considerations for electrodes to optimally activate the VN using endovascular electrodes in the IJV, as well as technologies to determine the position of the VN with respect to the IJV.

### Endovascular VNS requires precise electrode placement

We observed that one of the electrodes around the circumference of the vein wall resulted in lower current thresholds for stimulation evoked effects when stimulation was applied in monopolar configuration (Figure 3). This suggests that an endovascular VNS system would require accurate placement of the electrode within the vein such that the electrode is as close to the nerve as possible to both minimize the amount of stimulation energy required to activate the nerve and limit activation of other structures. There are a multitude of electrically excitable structures near the course of the VN such as the laryngeal branches of the VN, the phrenic nerve, the sympathetic trunk, hypoglossal nerve, accessory nerve, cardiac branches of the VN, and others. Therefore, methodology to ensure that a stent holding a small number of electrodes that are facing the nerve of interest, and away from other structures, during implantation will be critical.

We showed one such methodology that utilizes existing fluoroscopy and ultrasound tools are already used to implant endovascular devices, (Figure 5). Ultrasound is used to determine the orientation of the vein and nerve, ultrasound is used to guide a needle to place a fiducial near the VN, and fluoroscopy is used to guide the endovascular electrode to the location of the fiducial. The use of ultrasound to determine VN position relative to the carotid artery and IJV in humans was previously demonstrated [32,26]. Intravascular ultrasound has been demonstrated to be able to locate nerves through the renal artery [28], and thus may be a promising alternative or additional avenue to help locate the VN through the vein wall.

Another consideration for avoiding activation of nearby structures is to use bipolar electrode configurations. The strength of an electric field created at an electrode in bipolar configuration is known to decrease in proportion to 1/r^2^ where r is the distance from the electrode, while the electric field for a point source – a monopole – decreases in proportion to 1/r [24]. As expected, electrodes stimulating in bipolar configuration for this study showed increased stimulation thresholds compared to monopolar configuration (Figure 4). Therefore, bipolar configuration stimulation might be used to constrain the electric field and avoid activating nearby structures.

### Thresholds for nerve activation are higher for endovascular stimulation than cuff stimulation

VNS delivered from within the nearby IJV required more energy to activate the same nerve fiber types compared to a surgically placed electrode cuff using the same animal model and experimental preparation (RL-mediated EMG 1080uA endovascular vs 300uA surgical [14], and Aα CAP 860uA endovascular vs 300uA surgical [14]). An important consideration for the thresholds observed in this study is that the only method to optimize electrode position after placement via fluoroscopy was analysis of ENG and EMG stimulation evoked responses in the operating room. Therefore, we had limited time in choosing a position before performing the numerous stimulation parameters at the many electrode configurations tested in each animal. Thus, a more optimal electrode position may have been available in each animal but was not found.

The VN in swine and humans has been demonstrated to have complex cross-sectional morphology that changes along the length of the nerve, and thus could be expected to have locations that are better or worse for some clinically desired outcomes [10,11,27]. Indeed, studies stimulating the VN at cuff electrodes in swine and sheep have demonstrated that stimulating at different points along and around the nerve results in different evoked responses [12,13]. Likewise, studies of carotid sheath anatomy have demonstrated that the orientation of the VN relative to the IJV and carotid sheath are variable between patients [32]. Therefore, stimulation positions along the length of the nerve are likely to be dramatically different due to differences in nerve composition, distance between the nerve and vein, and tissue thicknesses. This anatomical variability strengthens the need for methodologies like ultrasound and fluoroscopy to be developed further for optimizing electrode position in the vein relative to the nerve. Sophisticated analysis of carotid sheath anatomy was not the focus of this initial study, but the histology from one animal demonstrates all of the above considerations of anatomical variability listed (Supplementary Figure 7). A future study focused on anatomy could be used to estimate likelihood of anatomy conducive to endovascular stimulation of the VN, as well as be used to seed computational models of endovascular stimulation (see below section) [25,31].

Even with optimization of electrode placement, one might assume that endovascular stimulation will never have stimulation effect thresholds as low as surgical placed electrodes given the distance and layers of tissue that impede endovascular stimulation. Two known physiological responses to device implants may benefit endovascular stimulation in a chronically implanted system. First, since there is no surgical cutdown, there will be no fibrous encapsulation of the nerve which may hinder cuff stimulation. Second, Oxley et al [20] have demonstrated that stenosis of their stent electrodes appears to improve signal to noise ratio over time, and could therefore be expected to perhaps form an insulation layer of endovascular stimulation electrodes from the blood and improve stimulation efficiency over time. Finally, despite increased stimulation effect thresholds, endovascular VNS might benefit from placement of the electrode in locations where certain fibers have already branched out of the nerve – such as locations in the thorax where the fibers of the recurrent laryngeal have already branched out of the VN trunk (Supplementary Figure 7).

With increased thresholds, electrode charge safety limits and battery capacity will also require additional evaluation. Stimulation at electrodes has been demonstrated to cause nerve and brain tissue damage depending on the charge density at the electrode [29,30], which is dependent on the size of the electrode and the current applied. Given that additional current is required to stimulate the nerve, additional attention will have to be paid to the size of the endovascular electrodes. Likewise, how stimulation through the vein wall or being in contact with highly conductive blood will affect the amount of charge density required to damage the tissue will require additional studies. Finally, assuming endovascular stimulation will require more current as surgical VNS, the lifetime and charging cycle of implantable pulse generator batteries will need to be considered in a cost-benefit analysis of a future system.

### Future directions leveraging computational modeling

As mentioned above, another important avenue for optimizing endovascular VNS is computational modeling. The results of this study were partially predicted by a computational model of endovascular VNS, where electrodes that were positioned on the edge of the vein closest to the nerve had lower thresholds than those that were offset [25]. Additional studies investigating vagotopy of the nerve [10,31], differences in distance between the vein and nerve, differences in tissue thicknesses, changes in endovascular electrode vessel wall integration [20] over time, and more using computational models could help estimate the best case scenario for endovascular VNS. Those models could also be used to suggest landmarks for imaging methodologies – such as ultrasound and fluoroscopy as we have suggested – in guiding optimal electrode location.

### Justification for using Aα CAP and EMG as output measures

The stimulation evoked responses we measured here might be considered to be only representative side effects of VNS. However, the responses we measured were demonstrated to be caused by activation of nerve fibers within the VN (Figure 2). Our goal in this study was to compare the relative thresholds for VN nerve fiber activation in response to different positions of electrodes in the nearby IJV, and thus the type of nerve fiber used for the comparison does not matter.

### Limitations

The Boston Scientific Orion cannot be used in humans chronically, therefore a custom electrode would need to be developed and optimized. The Orion fills much of the IJV in swine and may be blocking blood flow such that electrical shunting is not being appropriately modeled. The human IJV is approximately 3 times larger in diameter, therefore electrical shunting into the blood would likely have a larger effect, and electrode location will require greater precision.

## Conclusion

VNS used clinically is substantially limited by contraction of the neck muscles caused by activation of low threshold motor nerve fibers in the cervical VN. Here we investigated endovascular electrical stimulation within the IJV aimed at activation of the VN, which may allow for placement of a stimulation electrode at locations along the vagus that do not hold motor nerve fibers. As a starting point, we investigated how electrode position and configuration affect VN activation at the cervical VN, as we previously performed with the clinical helical cuff [14]. We found that optimal activation of the VN via endovascular stimulation required precise placement of the stimulation electrode around the circumference of the IJV. Bipolar configuration of multiple electrodes within the vein showed increased thresholds, but depended on the distance between the two bipole contacts. Position of endovascular stimulation electrodes with respect to the VN was verified via postmortem dissection and histology, which will not be possible for human patients. Thus, we also demonstrate methodologies via ultrasound and cone beam CT to determine the spatial relationship of the IJV and VN non-invasively.

This work establishes the feasibility of endovascular VNS with verification of nerve activation performed such that responses can be compared to a nearly identical demonstration using the clinically deployed helical cuff electrode [14]. While stimulation efficiency via endovascular stimulation is lower than using the helical cuff electrode, the slight differences in position of the endovascular electrode that cause large differences in threshold indicate that additional studies on how to target close anatomical configurations of the vein and nerve are warranted. Likewise, studies on anatomical variability of the VN and nearby veins would be useful to estimate best and worst case stimulation efficiency via computational modeling.

