## Supplementary Figures for "Vagus nerve stimulation using an endovascular electrode array"

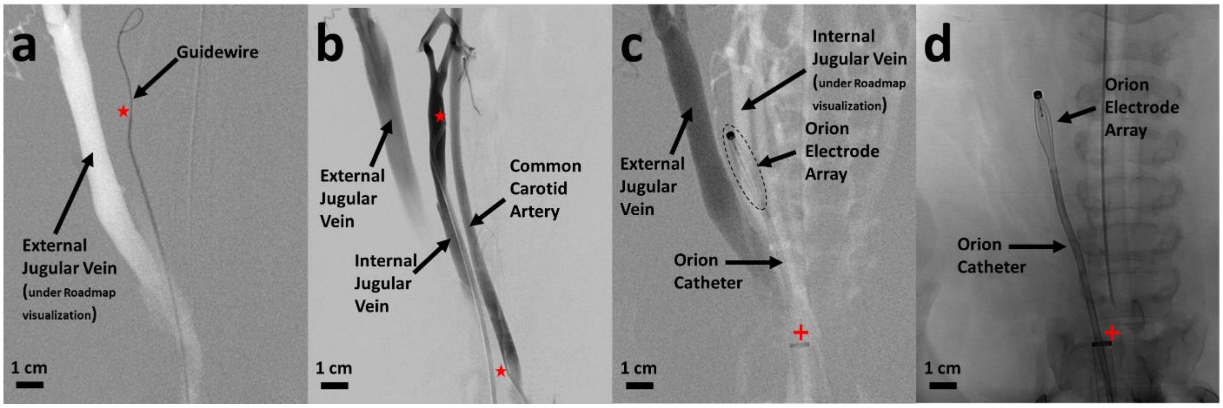


**Supplementary Figure 1:** Verification of electrode position within the internal jugular vein via series of angiograms/venograms. **a)** A retrograde roadmap venogram is produced during injection of Omnipaque contrast into the ear vein, which empties into the external jugular vein. The hyperintense vessel is therefore assumed to be the external jugular vein, and the guidewire (0.035” diameter microwire) within a 5Fr vertebral sheath (tip marked with red star) is placed medially to where the internal jugular vein is assumed to be positioned. The fluroscopic image shown was then captured. **b)** The guidewire is removed. The fluroscopic image shown was captured during simultaneous injection of contrast into the ear vein (catheter outside image window, to highlight the external jugular vein), injection of contrast into the 5Fr vertebral catheter from **a** (tip marked with red star, to highlight the internal jugular vein), and injection of contrast into the 5Fr sofia catheter (tip marked with red star, to highlight the common carotid artery). **c)** A roadmap visualization is produced during injection of contrast into the 5Fr sofia catheter (to highlight the internal jugular vein). The 5Fr vertebral catheter is removed, then replaced with an 18Fr sheath (tip marked with red cross). The Orion catheter is placed inside the 18Fr sheath and positioned via roadmap visualization. The electrode array on the Orion catheter is highlighted with a dotted oval. The fluroscopic image shown was then captured during injection of contrast into the ear vein to demonstrate that the Orion is outside the external jugular vein and is consistent with the position of the internal jugular vein roadmap visualization. **d)** The fluroscopic image shown was captured without injection of contrast and the Orion in final position before starting stimulation and recording experiments.

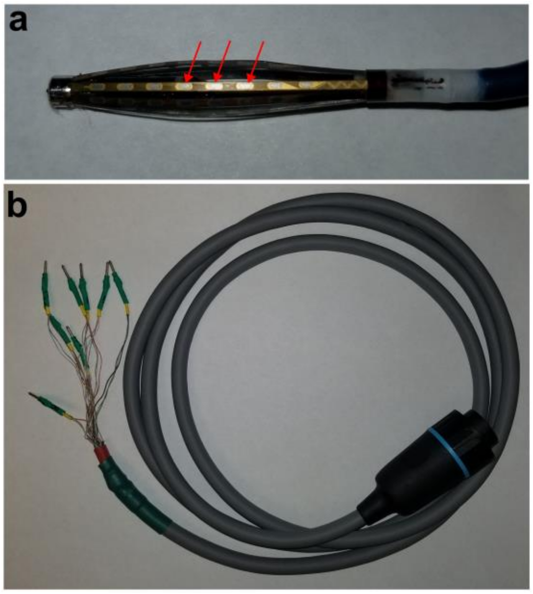


**Supplementary Figure 2:** Modification of Orion Umbilical Cable to connect multiple contacts on Orion catheter to increase electrode surface area. **a)** Photo of the Orion, showing multiple electrode contacts on one strut. Arrows show the three contacts that were combined – via modification of the umbilical cable in **b** – to form the electrode contact used for the experiments in this study. This process was repeated for each of the 8 Orion struts to form a ring of 8 electrodes. **b)** Photo of the modified umbilical cable. One of the proprietary connectors was cut off and individual lead wires were isolated. A multimeter was used to map each of the wires to individual contacts on the Orion. The wires leading to each set of three contacts were soldered together and fitted with a gold pin that was connected to the stimulator during experiments for each contact.

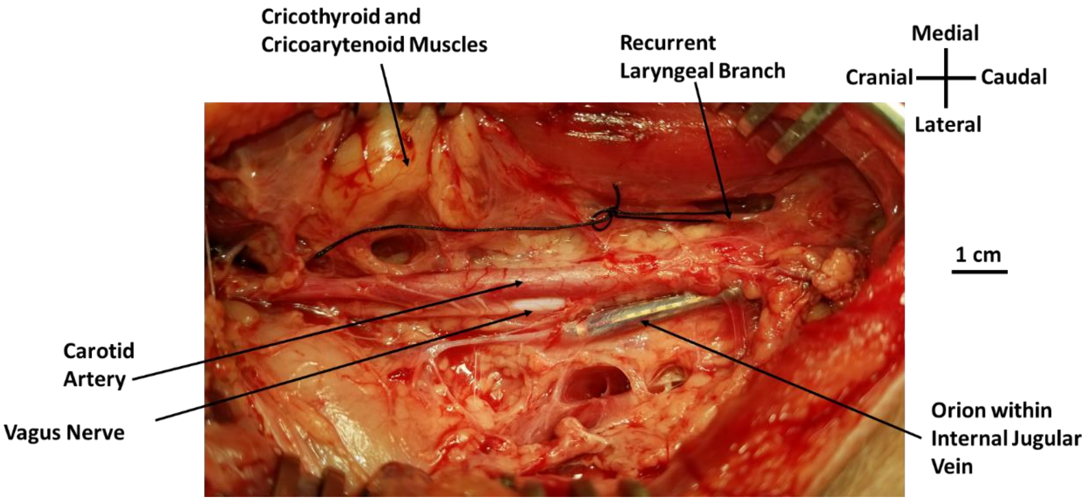


**Supplementary Figure 3:** Postmortem dissection to determine location of the stimulation electrode with respect to the vagus nerve. After experiments stimulating the nerve using the Orion at a location that was fully intact with no dissection, careful dissection was performed down to the level of the carotid sheath. Markers placed on the Orion were used to determine position of the Orion with respect to the vagus nerve, which is possible given the transparency of the vein.

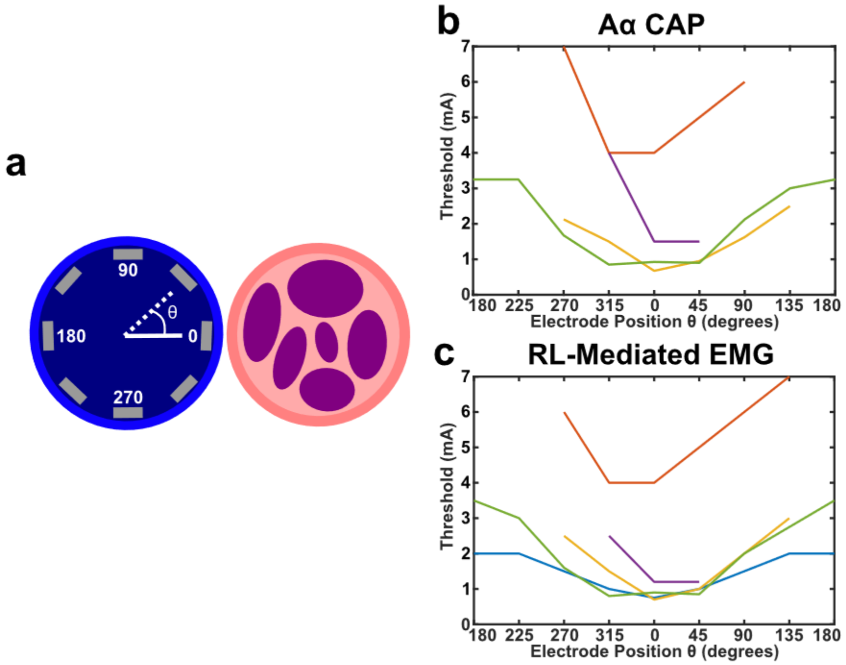


**Supplementary Figure 4:** Summary of thresholds in response to monopolar stimulation around the vein circumference across animals. **a)** Cartoon shows the internal jugular vein as the blue circle (left), vagus nerve trunk as the pink circle (right), and the 8 electrode contacts as the gray rectangles. Position of stimulation electrode contact numbers were verified with postmortem dissection, and data from each animal was binned such that the electrode closest to the VN were labeled as θ = 0. **b)** Mean thresholds for activation of the Aα CAP in response to monopolar stimulation at each of the 8 contacts in all animals (n = 4 pigs, different colored lines are different animals). **c)** Same as b, but for RL-mediated EMG (n = 5 pigs, different colored lines are different animals). For electrode positions in **b** and **c** that do not show data points, no response was observed up to 5.5 mA for the blue, green, yellow, and purple animals, and no response was observed up to 7 mA for the red animal potentially indicating a higher, but untested threshold.

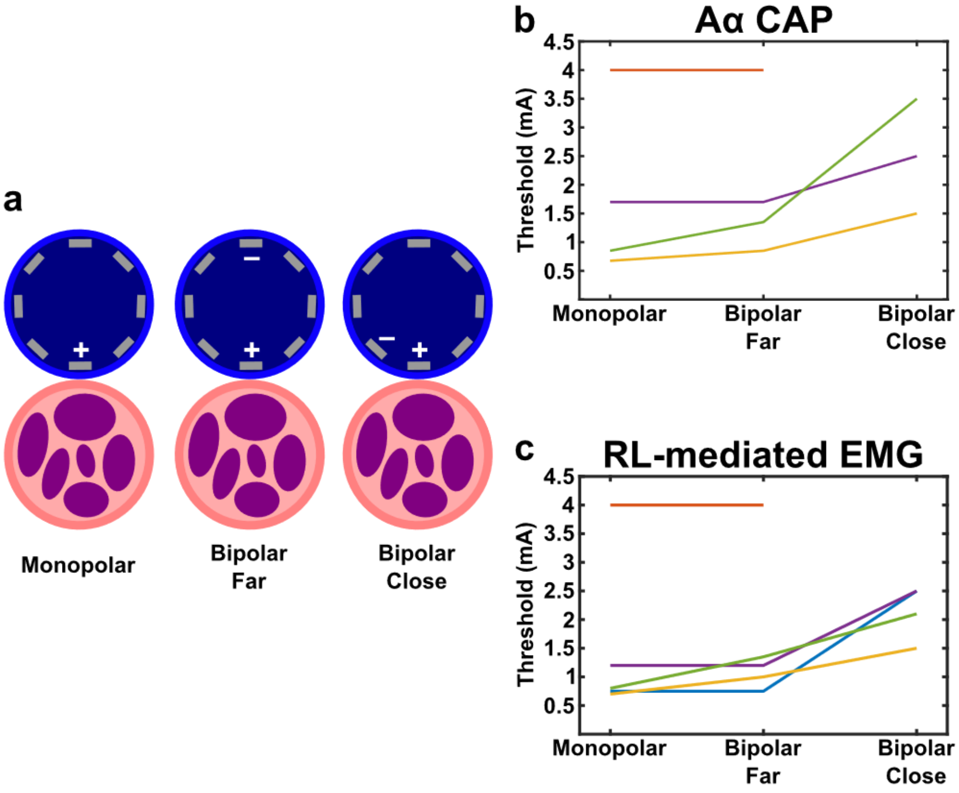


**Supplementary Figure 5:** Monopolar vs bipolar stimulation configurations across animals. **a)** Cartoons of vagus nerve (bottom circle) and internal jugular vein (top circle) showing the position of the electrodes for each monopolar and bipolar electrode configuration. Left cartoon is monopolar where the electrode closest to the nerve is being stimulated (+) and the return electrode is a needle in the left forelimb of the pig. Middle cartoon is bipolar "far" where the stimulated electrode (+) has a return at the electrode furthest away marked (-). Right cartoon is bipolar "close" where the stimulated electrode (+) has a return at the nearest electrode marked (-). **b)** Threshold for the Aα CAP in response to each configuration in all animals (n = 4 pigs, different colored lines are different animals). **c)** Threshold for the RL-mediated EMG in response to each configuration in all animals (n = 5 pigs, different colored lines are different animals). The animal with no response to bipolar close was stimulated up to 7 mA in that configuration and showed no response potentially indicating a higher, but untested threshold.

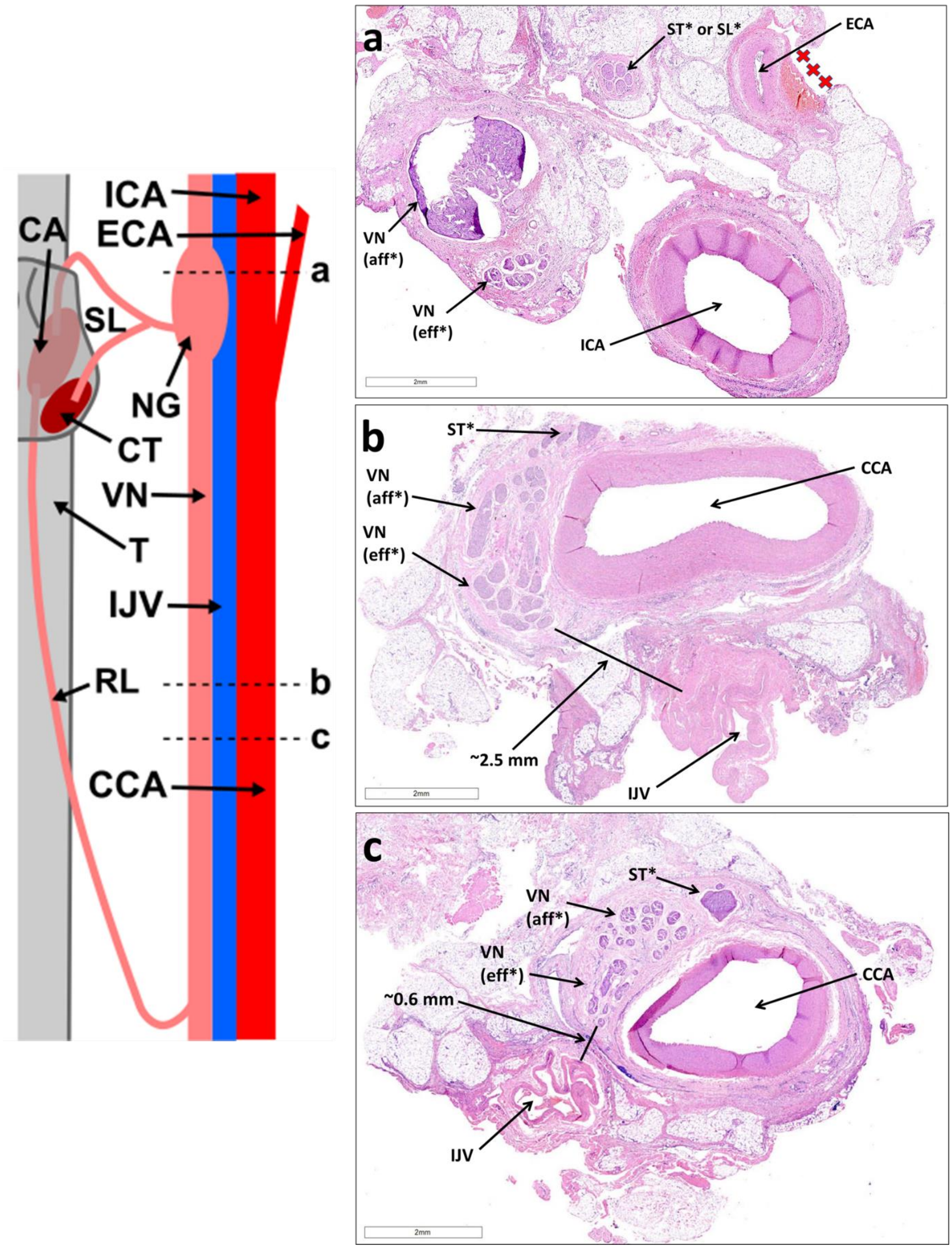
ICA
ECA
CA
SL
NG
CT
VN
T
IJV
RL
CCA
a
b
c
a
ST* or SL*
ECA
VN
(aff*)
VN
(eff*)
ICA
2mm
b
ST*
VN
(aff*)
VN
(eff*)
CCA
~2.5 mm
IJV
2mm
c
ST*
VN
(aff*)
VN
(eff*)
~0.6 mm
CCA
IJV
2mm

**Supplementary Figure 6:** Histology of the carotid sheath from one pig. **Left)** Cartoon of carotid sheath and surrounding anatomy. Dotted horizontal lines labeled with lowercase letters refer to where histology slides on right were approximately obtained. **Right)** Histology slides stained with hematoxylin and eosin (H&E). Relevant anatomy is labeled. Labels marked with an asterisk (*) indicate assumptions made by based on findings of Settell et al 2020 where internal vagus nerve (VN) organization of afferent (aff) and efferent (eff) fibers could be traced based on the position of the pseudounipolar cells of the nodose ganglion (NG) (see panel **a**, marked VN (aff)). That study also demonstrated that the sympathetic trunk (ST) may merge with the VN, thus the distinct third grouping of fascicles (or mode) near the VN might be the ST (marked with asterisk to highlight assumptions made). The Orion electrode array for this animal was placed near the histology shown in panels **b** and **c**. Note the 4x difference in distance between the VN and internal jugular vein (IJV) for panels **b** and **c** (2.5 mm vs 0.6 mm, respectively). Red crosses in **a** highlight hemorrhage likely associated with surgical cutdown for placement of longitudinal intrafascicular electrodes for nerve recording. Signs of hemorrhage could not be found for locations near the placement of the Orion electrode. Vagus nerve (VN), internal jugular vein (IJV), common carotid artery (CCA), internal carotid artery (ICA), external carotid artery (ECA), nodose ganglion (NG), trachea (T), recurrent laryngeal (RL), superior laryngeal (SL), cricoarytenoid muscle (CA), cricothyroid  muscle (CT), sympathetic trunk (ST), afferent nerve fibers (aff), efferent nerve fibers (eff).

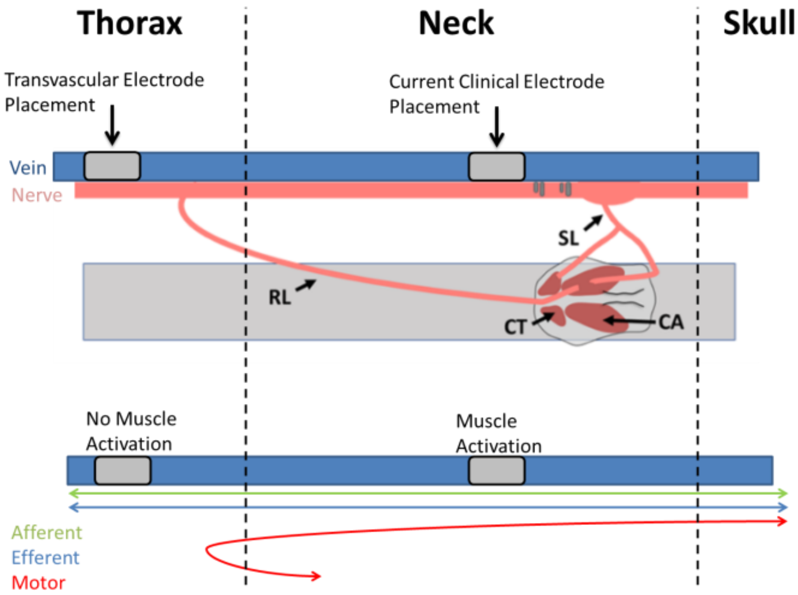


**Supplementary Figure 7:** Envisioned methodology to avoid activation of neck muscles using vagus nerve stimulation at an endovascular electrode. Top cartoon shows relevant anatomy and bottom cartoon shows wiring diagram of vagus nerve fibers. The surgically placed electrode in the clinic are be placed in the cervical region as the surgery is much less invasive than a craniotomy or thoracotomy. The fibers of the recurrent laryngeal nerve are located within the vagus trunk at all points along the cranial/caudal axis of the cervical vagus. Using an endovascular approach, the electrodes could be placed caudal to the recurrent laryngeal branching point on the vagus nerve in the thorax and activation of the motor efferents could be avoided.
